# An alternative interpretation of the slow KaiB-KaiC binding of the cyanobacterial clock proteins

**DOI:** 10.1101/2020.04.21.054270

**Authors:** Shin-ichi Koda, Shinji Saito

**Affiliations:** Department of Theoretical and Computational Molecular Science, Institute for Molecular Science, 38 Nishigo-Naka, Myodaiji, Okazaki, Aichi 444-8585, Japan; School of Physical Sciences, The Graduate University for Advanced Studies, 38 Nishigo-Naka, Myodaiji, Okazaki, Aichi 444-8585, Japan

## Abstract

The biological clock of cyanobacteria is composed of three proteins, KaiA, KaiB, and KaiC. The KaiB-KaiC binding brings the slowness into the system, which is essential for the long period of the circadian rhythm. However, there is no consensus as to the origin of the slowness due to the pre-binding conformational transition of either KaiB or KaiC. In this study, we propose a simple KaiB-KaiC binding scheme in a hexameric form with an attractive interaction between adjacent bound KaiB monomers, which is independent of KaiB’s conformational change. We then show that the present scheme can explain several important experimental results on the binding, including that used as evidence for the slow conformational transition of KaiB. The present result thus indicates that the slowness arises from KaiC rather than KaiB.

## Introduction

Circadian clocks are endogenous timing systems embedded in most living organisms and enable the organisms to adjust their biological activities to daily changes in the environment. Cyanobacteria are known as the simplest organisms that possess the circadian clock, which is composed of three proteins, KaiA, KaiB, and KaiC^1^. Since this circadian rhythm can be reconstituted in vitro just by mixing the three proteins with ATP^2^, this KaiABC oscillator has attracted much interest in elucidating the molecular mechanism of circadian rhythm.

KaiC forms a homohexamer with two ring-shaped domains, C1 and C2^3, 4^. In the presence of KaiA and KaiB, cyclic phosphorylation and dephosphorylation of two residues near the ATP binding site in C2, Ser431 and Thr432, occurs in the following order: ST → SpT → pSpT → pST → ST^5, 6^, where S/pS and T/pT are the unphosphorylated/phosphorylated states of Ser431 and Thr432, respectively. In this cycle, KaiA promotes the phosphorylation reaction^7, 8^ by acting on the C-terminal tail of C2^9–12^. KaiB, on the other hand, inhibits KaiA activity^13, 14^, which results in the dephosphorylation of C2. Specifically, after adequate phosphorylation of C2, KaiB binds to C1 and strongly sequesters KaiA from C2 by forming the C1-KaiB-KaiA complex^15^, which has recently been observed directly^16, 17.^

Several experimental studies have shown that the KaiB binding to C1 is slow, i.e. there exists a delay between the phosphorylation of C2 and the binding^5, 6, 18^. Owing to this delay, KaiC can be deeply phosphorylated, and hence the phosphorylation oscillation becomes robust^19, 20^. Yet, there has been a debate on the origin of the slowness of the KaiB-KaiC binding. An experiment by fluorescence anisotropy, which we refer to as Chang’s experiment, has shown that an initial rapid KaiB-KaiC binding is followed by a slow binding at a protein concentration of diluted KaiB (0.05 *µ*M of KaiB and 10.0 *µ*M of KaiC)^21^. This study has further shown that both the initial rapid and the subsequent slow bindings can be explained in a unified manner by assuming a conformational-selection binding scheme, i.e. a slow pre-binding conformational transition of KaiB dominates the binding. On the other hand, another experiment by fluorescence spectroscopy with tryptophan mutagenesis, which we refer to as Mukaiyama’s experiment, has shown that the rate of the KaiB-KaiC binding decreases with increasing the concentration of KaiB^22^. With a kinetic simulation based on a binding scheme considering pre-binding conformational transitions of both KaiB and KaiC, the study has concluded that the slowness of the binding arises from KaiC. In contrast to Chang’s experiment at only one protein concentration^21^, Mukaiyama’s experiment has measured the protein concentration dependence of the binding rate^22^, which generally provides a diagnostic clue to determine the slow conformational transition involved in a binding process^23^. Thus, it is conceivable that the conformational transition of KaiC is slower than KaiB. However, Mukaiyama’s experiment^22^ can explain only the initial rapid binding in Chang’s experiment^21^ because Mukaiyama’s experiment^22^ shows a high binding rate when KaiB is dilute. Therefore, an explanation that satisfies both of the two experiments, in particular, the slow binding observed in Chang’s experiment^21^, is needed.

In this article, we propose an alternative binding scheme that can explain both the initial rapid and the subsequent slow KaiB-KaiC binding observed in Chang’s experiment^21^. In contrast to the original conformational-selection scheme proposed in Chang’s experiment^21^, the present scheme works even when the conformational transition of KaiB is arbitrarily fast, which is thus also consistent with the conclusion of Mukaiyama’s experiment^22^. The key points of the present scheme are three-fold: the low concentration of KaiB^21^, the hexameric form of KaiC, and the attractive adjacent KaiB-KaiB interaction in the KaiB-KaiC complex^17, 24^. In the next section, we show that the slowness of the binding in Chang’s experiment^21^ may arise from the low concentration of KaiB. Specifically, in the present scheme, the initial rapid and the subsequent slow bindings correspond to the bindings from C_6_B_0_ to C_6_B_1_ and from C_6_B_1_ to the other larger KaiB-KaiC complexes C_6_B_*n*_ (*n >* 1), respectively. Here, C_6_B_*n*_ represents the complex of a KaiC hexamer and *n* KaiB monomers. Note that, without any stabilization mechanism, the formation of the larger KaiB-KaiC complexes C_6_B_*n*_ (*n >* 1) seldom occurs at a low concentration of KaiB. In the present scheme, however, the larger KaiB-KaiC complexes are substantially formed due to the stabilization by the attractive adjacent KaiB-KaiB interaction.

## Results

### Design of binding scheme

In this study, we discuss the KaiB-KaiC binding with a binding scheme in which the hexameric form of KaiC is explicitly considered (Fig. 1). We distinguish all six binding sites on a KaiC hexamer and consider 2^6^ types of the KaiB-KaiC complex, i.e. whether KaiB is bound at each binding site (not explicitly shown in Fig. 1). The conformational transition of KaiB is assumed to be fast and is treated by the so-called rapid equilibrium approximation. Thus, the mutagenesis of KaiB to stabilize the binding-competent conformation^21^ is effectively represented by the increase in the association rate constant of the KaiB-KaiC binding. To focus on the binding in Chang’s experiment^21^, we ignore the conformational transition of KaiC and assume that all KaiC hexamers take their binding-competent conformational state.

**Figure 1.**
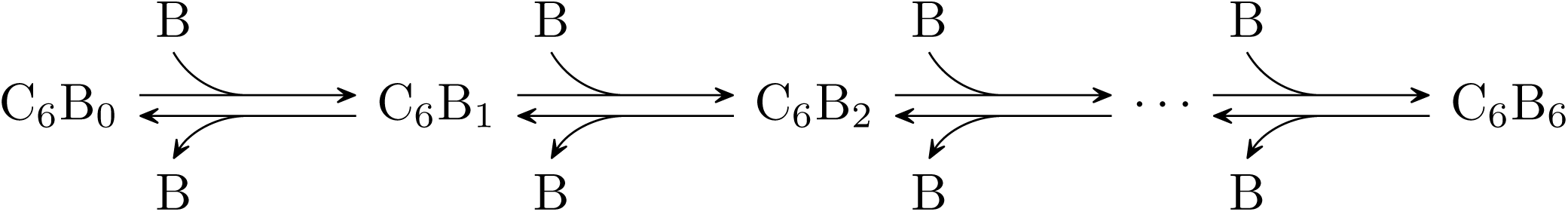
Present scheme for the KaiB-KaiC binding. C_6_B_*n*_ (*n* = 0, 1, *…*, 6) are the complexes of a KaiC hexamer and *n* KaiB monomers, and B represents a KaiB monomer.

We set the association and dissociation rate constants as follows. For simplicity, we assume that the association rate constant of a KaiB monomer to a KaiB-KaiC complex is always *k*_on_ (Fig. 2). On the other hand, we assume that the dissociation of a KaiB monomer from a KaiB-KaiC complex becomes slow when another KaiB monomer is bound at an adjacent binding site. This is because the attractive KaiB-KaiB interaction^17, 24^ may stabilize the complex. Specifically, the dissociation rate constant *k*_off_ is modeled as

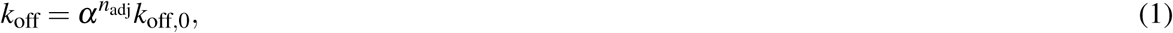

where *n*_adj_ is the number of adjacent KaiB monomers, *α* (*<* 1) is the factor representing the stabilization by the KaiB-KaiB interaction, and *k*_off,0_ is the rate constant without breaking the KaiB-KaiB interaction. Note that Eq. (1) satisfies the thermodynamic consistency, i.e. the detailed balance condition in the transitions among C_6_B_*n*_ (0 *≤ n ≤* 6). Some examples of the dissociation are shown in Fig. 2.

**Figure 2.**
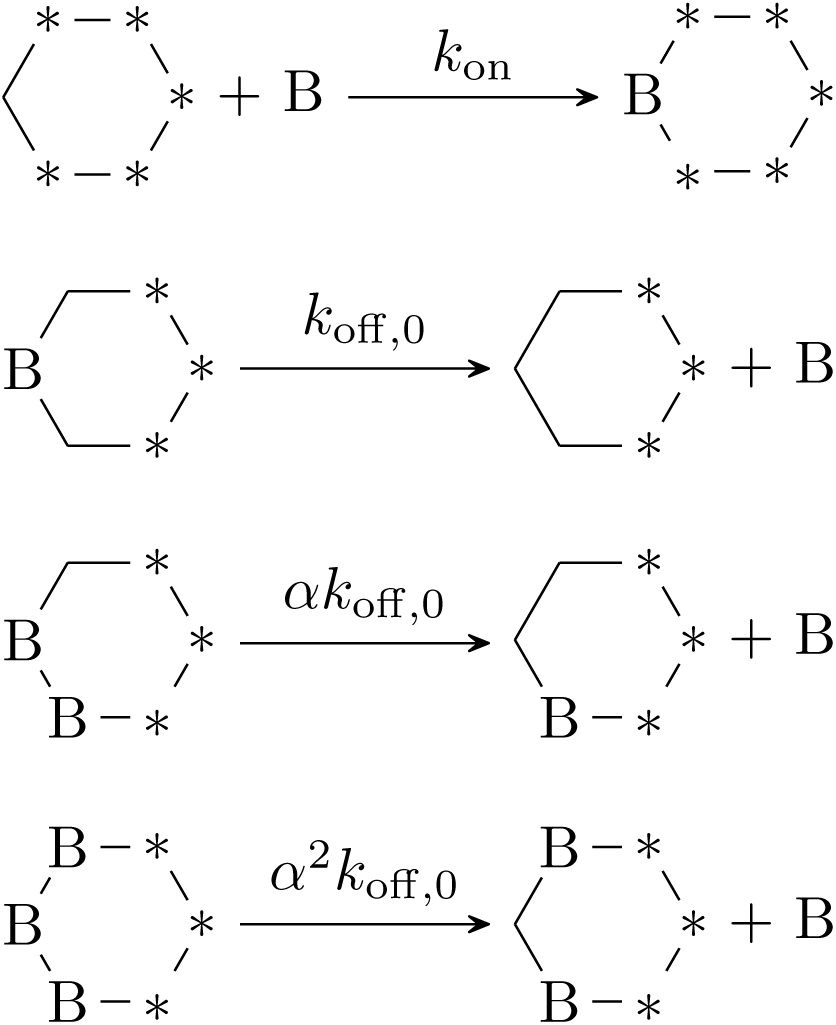
Association and dissociation of a KaiB monomer to/from a KaiB-KaiC complex. A hexagon represents a KaiC hexamer. Vertexes with and without B represent the occupied and unoccupied binding sites, respectively. An asterisk indicates that the corresponding binding site is either of the occupied or unoccupied sites. The first scheme represents the association. The association rate constant is independent of independent of the presence of B at other binding sites. The last three schemes summarize the dissociation. The dissociation rate constant depends on the number of adjacent KaiB monomers due to the KaiB-KaiB interaction.

### Origin of the slowness

We then show that the present scheme with appropriate parameter values can qualitatively reproduce the time course of the KaiB-KaiC binding in Chang’s experiment^21^. To make the following discussion clear, we first show a numerical result of the present scheme (Fig. 3A). Here, the protein concentrations are set to be the same as in Chang’s experiment^21^, i.e. the total concentrations of KaiB, B_tot_, and of KaiC, C_tot_, are 0.05 *µ*M and 10.0 *µ*M, respectively. We also set the other parameters as *k*_off,0_ = 200.0 s^−1^, *α* = 0.001, and *k*_on_ = 2.0, 5.0, 10.0 *µ*M^−1^s^−1^ so that the present scheme can reproduce the experimental results^21, 22, 24^. The result of the present scheme shown in Fig. 3A is consistent with the binding observed in Chang’s experiment^21^ in terms of the following three features. First, the present scheme can reproduce both the initial rapid and the subsequent slow bindings. Second, an increase in *k*_on_, which corresponds to the mutagenesis of KaiB to stabilize the binding-competent conformation, leads to the increases in the amount of bound KaiB both in the initial and the subsequent bindings. Lastly, the KaiB-KaiC binding is accelerated as *k*_on_ increases.

**Figure 3.**
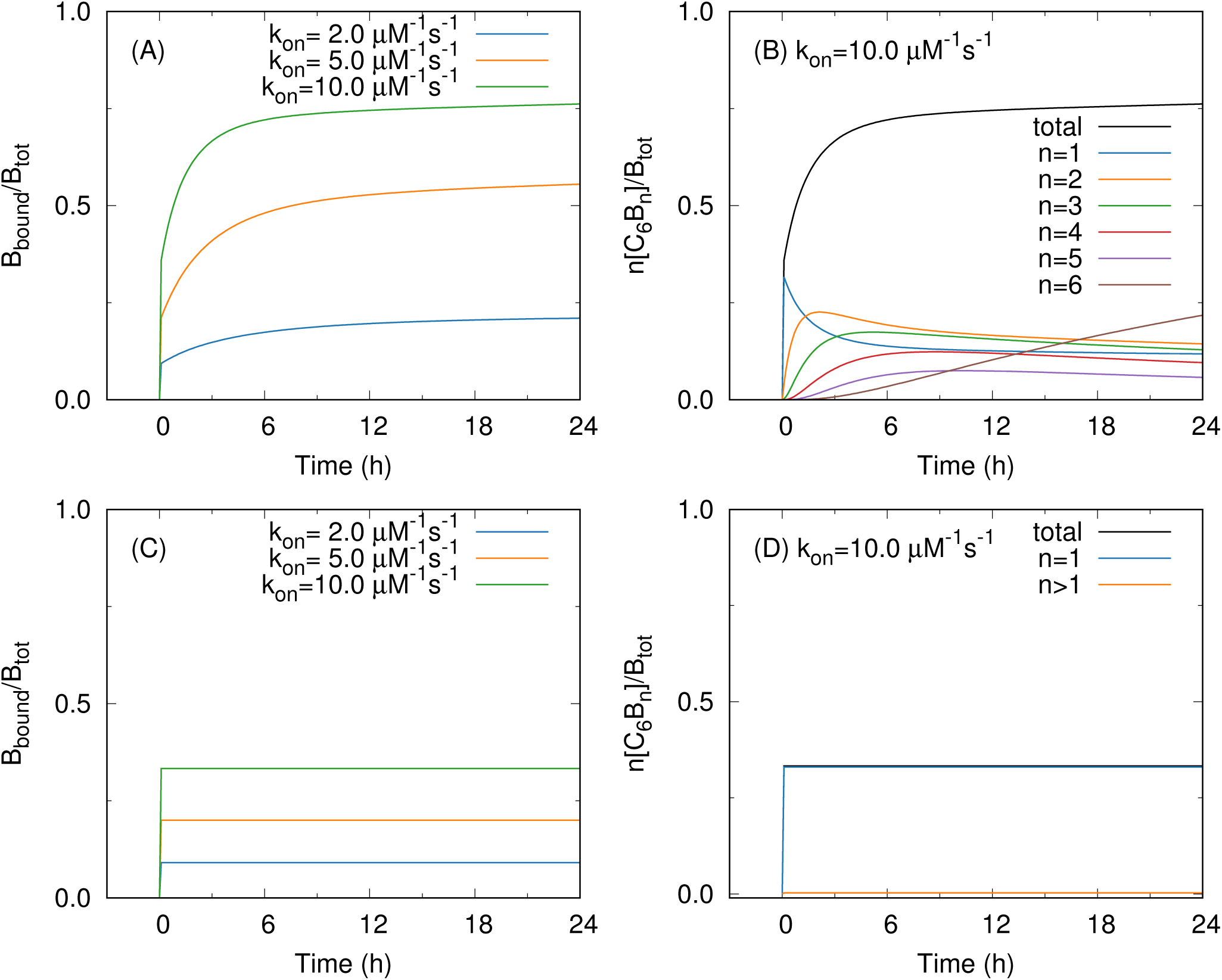
Time courses of bound KaiB at the protein concentrations B_tot_ = 0.05 *µ*M and C_tot_ = 10.0 *µ*M with the dissociation rate constant *k*_off,0_ = 200.0 s^−1^. (A) and (C) show *k*_on_ dependence of bound KaiB with *k*_on_ = 2.0, 5.0, 10.0 *µ*M^−1^s^−1^. (B) and (D) show bound KaiB in C_6_B_*n*_ (*n* = 1, *…*, 6) when *k*_on_ = 10.0 *µ*M^−1^s^−1^. The adjacent KaiB-KaiB interaction is considered with *α* = 0.001 in (A) and (B), whereas it is ignored with *α* = 1.0 in (C) and (D).

Next, we explain why the present scheme can reproduce the KaiB-KaiC binding in Chang’s experiment^21^. The initial rapid and the subsequent slow binding can be explained by the low concentration of KaiB (B_tot_ = 0.05 *µ*M, C_tot_ = 10.0 *µ*M) and the hexameric form of KaiC. The rate of the KaiB-KaiC binding at an unoccupied binding site of C_6_B_*n*_ (*n* = 0, 1, *…*, 5), *v*_*n*_, is given by

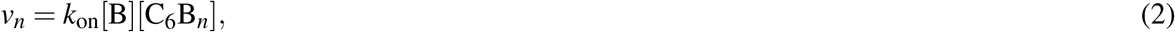

where [B] and [C_6_B_*n*_] are the concentrations of unbound KaiB and C_6_B_*n*_, respectively. In the case of the low concentration of KaiB, most KaiC hexamers bind no KaiB monomer because the concentration of KaiC is much larger than that of KaiB. Thus, [C_6_B_0_] can be approximated as

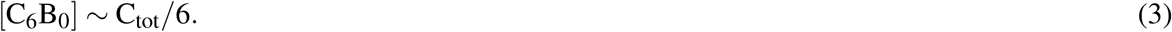

On the other hand, [C_6_B_*n*_] (*n ≥* 1) never exceeds B_tot_, i.e.

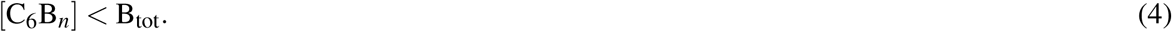

Therefore, we obtain

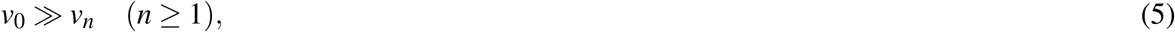

which indicates that the initial binding from C_6_B_0_ to C_6_B_1_ is much faster than the subsequent binding from C_6_B_*n*_ to C_6_B_*n*+1_ (*n ≥* 1). This is why the subsequent binding is much slower than the initial binding in the present scheme. Figure 3B indeed shows that the initial rapid binding is attributed to the binding from C_6_B_0_ to C_6_B_1_ and the subsequent slow binding to the binding from C_6_B_1_ to the larger KaiB-KaiC complexes.

In the present scheme, we consider the KaiB-KaiB interaction for the stabilization of larger KaiB-KaiC complexes^24^. Without the KaiB-KaiB interaction, the KaiB-KaiC binding at a binding site becomes independent of other binding sites. Thus the distribution of [C_6_B_*n*_] (0 *≤ n ≤* 6) at equilibrium obeys a binominal distribution. In this case, the amount of larger KaiB-KaiC complexes becomes much smaller than [C_6_B_1_] when KaiB is dilute, which means that the binding from C_6_B_*n*_ to C_6_B_*n*+1_ (*n ≥* 1) seldom occurs. The numerical results without the stabilization by the KaiB-KaiB interaction, in which *α* is set to be 1 and the other parameter values are the same as those used in Figs. 3A and B (Figs. 3C and D), indeed show that the slow binding to the larger KaiB-KaiC complexes is absent in this case. On the other hand, with the stabilization by the KaiB-KaiB interaction, the concentration of a complex in C_6_B_*n*_ with *m* KaiB-KaiB interfaces at equilibrium becomes

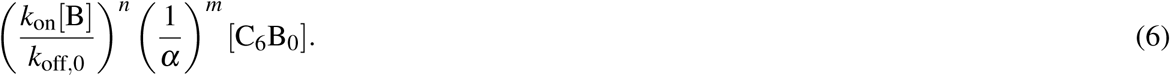

Therefore, with appropriately small *α*, [C_6_B_*n*_] (*n ≥* 2) becomes comparable to [C_6_B_1_] at equilibrium, as we have already seen in Figs. 3A and B. This is why the amount of the subsequent binding is comparable to the initial binding in the present scheme.

Equations (2) and (6) can also explain *k*_on_ dependence of the binding. Equation (2) indicates that the binding is accelerated as *k*_on_ increase, and Eq. (6) means that the amount of the binding increase with *k*_on_ both in the initial and subsequent bindings due to the equilibrium shift toward larger KaiB-KaiC complexes.

### KaiB-KaiC binding at other protein concentrations

In this subsection, we investigate the KaiB-KaiC binding at protein concentrations different from the low KaiB concentration. A recent experiment by mass spectrometry has shown that C_6_B_6_ is the most abundant in C_6_B_*n*_ (*n* = 1, 2, *…*, 6) six hours after mixing KaiB with a phospho-mimic mutant of KaiC, even when B_tot_ is smaller than C_tot_ (B_tot_*/*C_tot_ = 0.25)^24^. The present scheme can describe this experimental result (Fig. 4A) because the ring-shaped alignment of KaiB monomers in C_6_B_6_ acquires the KaiB-KaiB interaction more efficiently than C_6_B_*n*_ (*n <* 6) as follows. When *n <* 6, the number of KaiB-KaiB interfaces in C_6_B_*n*_ is not more than *n*− 1. On the other hand, the number of interfaces is *n* when *n* = 6 because the bound KaiB monomers form a ring. Therefore, if the stabilization effect *α* is small enough to be comparable to *k*_on_[B]*/k*_off,0_ at equilibrium, i.e.

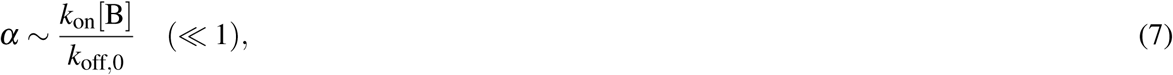

**Figure 4.**
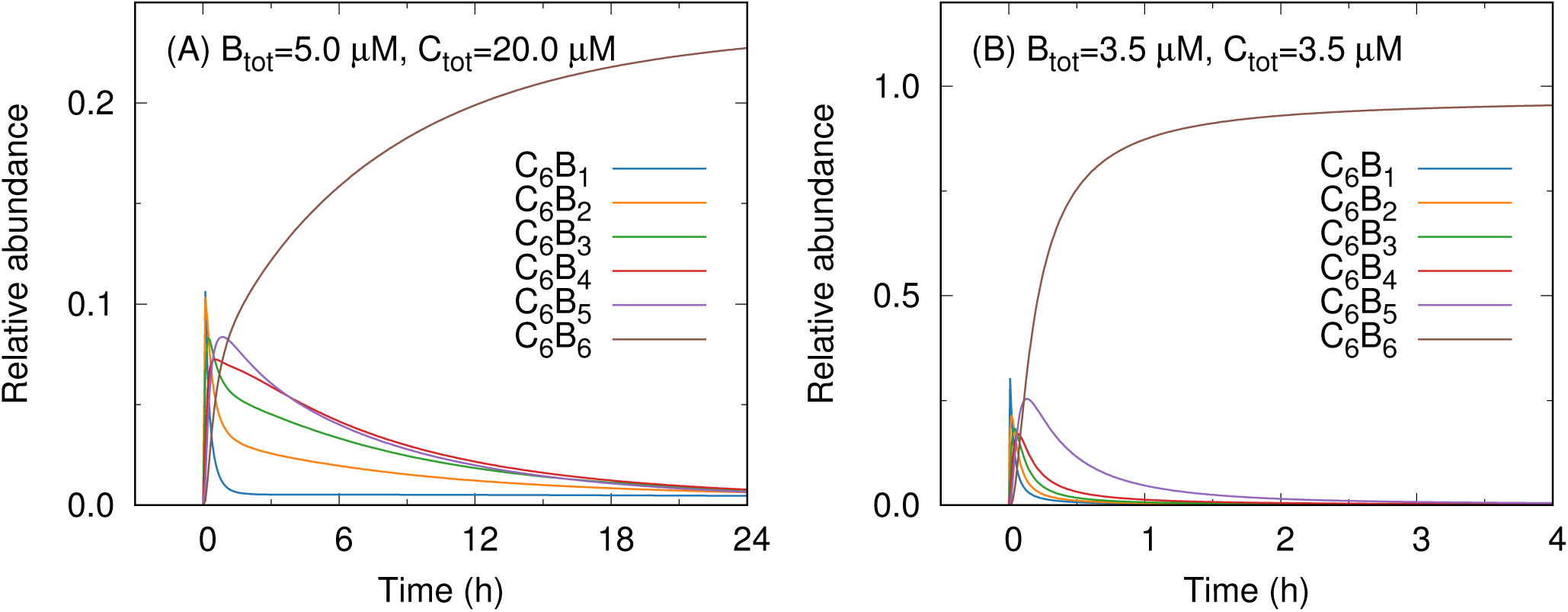
Relative abundances of the KaiB-KaiC complexes in the binding with *k*_on_ = 2.0 *µ*M^−1^s^−1^, *k*_off,0_ = 200.0 s^−1^, and *α* = 0.001 at the protein concentrations (A) B_tot_ = 5.0 *µ*M and C_tot_ = 20.0 *µ*M, and (B) B_tot_ = C_tot_ = 3.5 *µ*M.

Eq. (6) yields

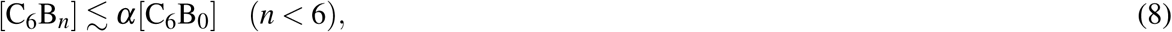

and

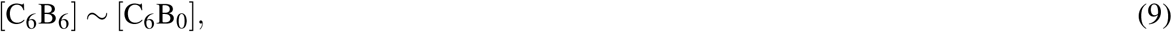

which indicate that [C_6_B_6_] is much larger than other [C_6_B_*n*_] (1 *≤ n <* 6).

Lastly, we investigate the KaiB-KaiC binding at the so-called standard condition of the KaiABC oscillator (B_tot_ = C_tot_ = 3.5 *µ*M). As Eq. (2) indicates that the binding is accelerated with increasing protein concentrations, the result of the present scheme exhibits a rapid binding with the half-life time *t*_1*/*2_ = 0.06 h at the standard condition (Fig. 4B). Although this half-life time is considerably shorter than that observed in Mukaiyama’s experiment^22^, the present result does not contradict with the experimental result. This is because the present scheme ignores the conformational transition of KaiC. If we incorporate a slow pre-binding conformational transition of KaiC into the present scheme in a conformational-selection manner, the binding rate could be regulated by KaiC as suggested in Mukaiyama’s experiment^22^. Rather, the present result (Fig. 4B) indicates that the slowness of the binding arising from protein concentration may be negligible at the standard condition compared to that from the conformational transition of KaiC, i.e. the KaiB-KaiC binding itself is rapid and may not be a rate-limiting process in the KaiABC oscillator around the standard condition. This result is also consistent with the experimental result that an equimolar increase in the concentrations of KaiA, KaiB, and KaiC unalters the phosphorylation oscillation^25^. If the binding process is so slow that it depends on the protein concentration, the change of the protein concentration would modulate the oscillation.

## Discussion

In this study, we proposed an alternative interpretation of the KaiB-KaiC binding based on a novel binding scheme, which is consistent with the three experimental results: the initial rapid and the subsequent slow binding at a low KaiB concentration^21^, the protein concentration dependence of the binding rate^22^, and the dominant formation of C_6_B_6_ even when B_tot_ *<* C_tot_^24^.To the best of our knowledge, these experiments have not been explained in a unified way before.

The binding scheme used in this study simply models the hexameric form of the KaiB-KaiC complex and the attractive adjacent KaiB-KaiB interaction in the complex. As the structural study has revealed a strong electrostatic interaction between two adjacent KaiB monomers in the complex^17^, it is reasonable to assume that the dissociation of a KaiB monomer from the complex becomes slow when the KaiB-KaiB interaction is formed. It should be noted that the parameter values used in the present study are determined by hand, i.e. without any numerical fitting to experimental data. Thus, the values themselves are of little importance. However, the framework of the present scheme is still of importance because it gives the possible comprehensive mechanism of the KaiB-KaiC binding summarized below.

We showed that the present scheme can qualitatively reproduce the initial rapid and the subsequent slow binding^21^ at a low KaiB concentration along the following scenario. When KaiB is much fewer than KaiC, the amount of the bare KaiC hexamer C_6_B_0_ substantially exceeds the KaiB-KaiC complexes C_6_B_*n*_ (*n ≥* 1). Thus, the initial binding from C_6_B_0_ to C_6_B_1_ becomes much faster than the subsequent binding form C_6_B_1_ to the larger complexes C_6_B_*n*_ (*n >* 1). Moreover, due to the stabilization by the adjacent KaiB-KaiB interaction, a comparable amount of the larger complex to C_6_B_1_ is produced. Note that this binding mechanism does not rely on the structural transition of KaiB, which indicates that the rate of the transition can be arbitrarily fast. In the present study, by assuming that the transition is very fast and effectively changing the association rate constant *k*_on_, we showed that the present scheme can reproduce the mutagenesis dependence of the binding in Chang’s experiment^21^. Therefore, the slow binding observed in Chang’s experiment^21^ does not necessarily mean that the structural transition of KaiB is slow.

The present scheme can further explain why C_6_B_6_ is dominantly produced even when B_tot_ *<* C_tot_^24^ This is because the ring-shaped alignment of bound KaiB monomers in C_6_B_6_ has an advantage in the formation of KaiB-KaiB interaction compared to other alignments consisting of one or more linear segments.

We also showed that, without the slow pre-binding conformational transition of KaiC suggested in Mukaiyama’s experiment^22^, the KaiB-KaiC binding may become rapid at the standard condition of the KaiABC oscillator i.e. B_tot_ = C_tot_ = 3.5 *µ*M (Fig. 4B). This means that the conformational transition of KaiC could be the slowest process in the whole KaiB-KaiC binding process. In this sense, the present scheme is consistent with the experimental study indicating that the slowness of the binding arises from KaiC^22^.

In summary, the present study indicates that the slowness arises from KaiC and that the conformational transition of KaiB is of less importance in the binding. This conclusion is also supported by a recent mathematical reaction model of the KaiABC oscillator, which shows that various experimental results can be reproduced without considering the conformational transition of KaiB^26^.

It must be noted, however, that the conformational transition of KaiB can still play an essential role in the KaiABC oscillator. Indeed, the mutagenesis of KaiB to stabilize the binding-competent state of KaiB results in arrhythmic phosphorylation of KaiC in the presence of both KaiA and KaiB^21^. Thus, further investigation of the functional role of the conformational transition of KaiB is still needed in the future.

## Methods

### Numerical calculation

The rate equations of the present binding scheme are integrated by the fourth-order Runge-Kutta method with the time step Δ*t* = 0.001 s.

## Data availability

All data generated in this study are available from the corresponding authors upon request.

## Acknowledgements

This work has been supported by JSPS KAKENHI, Grant Number JP18K14185 (SK) and JP16H02254 (SS), and the Indo-Japan bilateral collaboration program. The calculations were partially carried out on computers at the Research Center for Computational Science, Okazaki, Japan.

## Author contributions

S.K. and S.S. designed research. S.K. performed research. S.K. and S.S. analyzed data and wrote the paper.

## References

1. Ishiura, M. et al. Expression of a gene cluster kaiabc as a circadian feedback process in cyanobacteria. Science 281, 1519–1523, DOI: 10.1126/science.281.5382.1519 (1998).

2. Nakajima, M. et al. Reconstitution of circadian oscillation of cyanobacterial kaic phosphorylation in vitro. Science 308, 414–415, DOI: 10.1126/science.1108451 (2005).

3. Mori, T. et al. Circadian clock protein kaic forms atp-dependent hexameric rings and binds dna. Proc. Natl. Acad. Sci. 99, 17203–17208, DOI: 10.1073/pnas.262578499 (2002).

4. Hayashi, F. et al. Atp-induced hexameric ring structure of the cyanobacterial circadian clock protein kaic. Genes to Cells 8, 287–296, DOI: 10.1046/j.1365-2443.2003.00633.x (2003).

5. Rust, M. J., Markson, J. S., Lane, W. S., Fisher, D. S. & O’Shea, E. K. Ordered phosphorylation governs oscillation of a three-protein circadian clock. Science 318, 809–812, DOI: 10.1126/science.1148596 (2007).

6. Nishiwaki, T. et al. A sequential program of dual phosphorylation of kaic as a basis for circadian rhythm in cyanobacteria. The EMBO J. 26, 4029–4037, DOI: 10.1038/sj.emboj.7601832 (2007).

7. Iwasaki, H., Nishiwaki, T., Kitayama, Y., Nakajima, M. & Kondo, T. Kaia-stimulated kaic phosphorylation in circadian timing loops in cyanobacteria. Proc. Natl. Acad. Sci. 99, 15788–15793, DOI: 10.1073/pnas.222467299 (2002).

8. Nishiwaki, T. et al. Role of kaic phosphorylation in the circadian clock system of synechococcus elongatus pcc 7942. Proc. Natl. Acad. Sci. United States Am. 101, 13927–13932, DOI: 10.1073/pnas.0403906101 (2004).

9. Vakonakis, I. & LiWang, A. C. Structure of the c-terminal domain of the clock protein kaia in complex with a kaic-derived peptide: Implications for kaic regulation. Proc. Natl. Acad. Sci. United States Am. 101, 10925–10930, DOI: 10.1073/pnas.0403037101 (2004).

10. Pattanayek, R. et al. Visualizing a circadian clock protein crystal structure of kaic and functional insights. Mol. Cell 15, 375–388, DOI: 10.1016/j.molcel.2004.07.013 (2004).

11. Kim, Y.-I., Dong, G., Carruthers, C. W., Golden, S. S. & LiWang, A. The day/night switch in kaic, a central oscillator component of the circadian clock of cyanobacteria. Proc. Natl. Acad. Sci. 105, 12825–12830, DOI: 10.1073/pnas.0800526105 (2008).

12. Tseng, R. et al. Cooperative kaia–kaib–kaic interactions affect kaib/sasa competition in the circadian clock of cyanobacteria. J. Mol. Biol. 426, 389–402, DOI: 10.1016/j.jmb.2013.09.040 (2014).

13. Kitayama, Y., Iwasaki, H., Nishiwaki, T. & Kondo, T. Kaib functions as an attenuator of kaic phosphorylation in the cyanobacterial circadian clock system. The EMBO J. 22, 2127–2134, DOI: 10.1093/emboj/cdg212 (2003).

14. Xu, Y., Mori, T. & Johnson, C. H. Cyanobacterial circadian clockwork: roles of kaia, kaib and the kaibc promoter in regulating kaic. The EMBO J. 22, 2117–2126, DOI: 10.1093/emboj/cdg168 (2003).

15. Chang, Y.-G., Tseng, R., Kuo, N.-W. & LiWang, A. Rhythmic ring–ring stacking drives the circadian oscillator clockwise. Proc. Natl. Acad. Sci. 109, 16847–16851, DOI: 10.1073/pnas.1211508109 (2012).

16. Snijder, J. et al. Structures of the cyanobacterial circadian oscillator frozen in a fully assembled state. Science 355, 1181–1184, DOI: 10.1126/science.aag3218 (2017).

17. Tseng, R. et al. Structural basis of the day-night transition in a bacterial circadian clock. Science 355, 1174–1180, DOI: 10.1126/science.aag2516 (2017).

18. Qin, X. et al. Intermolecular associations determine the dynamics of the circadian KaiABC oscillator. Proc. Natl. Acad. Sci. 107, 14805–14810, DOI: 10.1073/pnas.1002119107 (2010).

19. Phong, C., Markson, J. S., Wilhoite, C. M. & Rust, M. J. Robust and tunable circadian rhythms from differentially sensitive catalytic domains. Proc. Natl. Acad. Sci. 110, 1124–1129, DOI: 10.1073/pnas.1212113110 (2013).

20. Paijmans, J., Lubensky, D. K. & Wolde, P. R. t. A thermodynamically consistent model of the post-translational kai circadian clock. PLOS Comput. Biol. 13, e1005415, DOI: 10.1371/journal.pcbi.1005415 (2017).

21. Chang, Y.-G. et al. A protein fold switch joins the circadian oscillator to clock output in cyanobacteria. Science 349, 324–328, DOI: 10.1126/science.1260031 (2015).

22. Mukaiyama, A. et al. Conformational rearrangements of the c1 ring in kaic measure the timing of assembly with kaib. Sci. Reports 8, 8803, DOI: 10.1038/s41598-018-27131-8 (2018).

23. Vogt, A. D. & Cera, E. D. Conformational Selection or Induced Fit? A Critical Appraisal of the Kinetic Mechanism. Biochemistry 51, 5894–5902, DOI: 10.1021/bi3006913 (2012).

24. Murakami, R. et al. Cooperative Binding of KaiB to the KaiC Hexamer Ensures Accurate Circadian Clock Oscillation in Cyanobacteria. Int. J. Mol. Sci. 20, 4550, DOI: 10.3390/ijms20184550 (2019).

25. Kageyama, H. et al. Cyanobacterial Circadian Pacemaker: Kai Protein Complex Dynamics in the KaiC Phosphorylation Cycle In Vitro. Mol. Cell 23, 161–171, DOI: 10.1016/j.molcel.2006.05.039 (2006).

26. Koda, S.-i. & Saito, S. Sufficiency of unidirectional allostery in KaiC in generating the cyanobacterial circadian rhythm. bioRxiv 2020.04.01.021055, DOI: 10.1101/2020.04.01.021055 (2020).

